## Supplementary Information for "Strand-specific single-cell methylomics reveals distinct modes of DNA demethylation dynamics during early mammalian development"

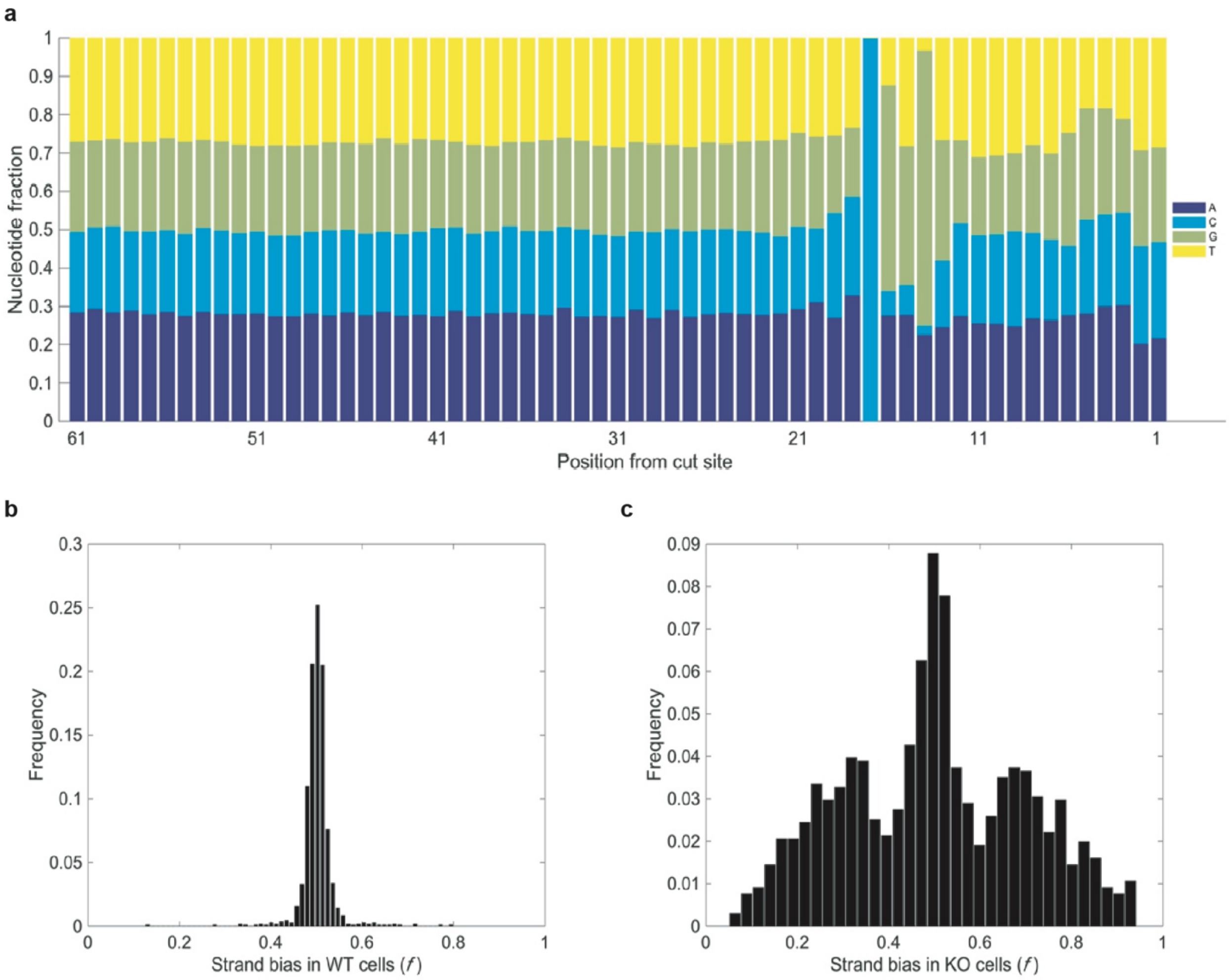


**Supplementary Figure 1 | Strand-specific detection of 5mC in single cells using scMspJI-seq.** (**a**) Panel shows the nucleotide composition that was observed downstream of the MspJI cut site. In agreement with previous reports, MspJI was found to cut gDNA 16 bp downstream of the cut site[1]. (**b**) Chromosomes in E14 cells show a tight strand bias distribution centered around 0.5. (**c**) CRISPR-cas9 mediated knockout of Dnmt1 in E14 cells results in a dramatic increase in the width of the strand bias distribution indicating loss of maintenance methylation and the ability of scMspJI-seq to quantify strand-specific 5mC in single cells.

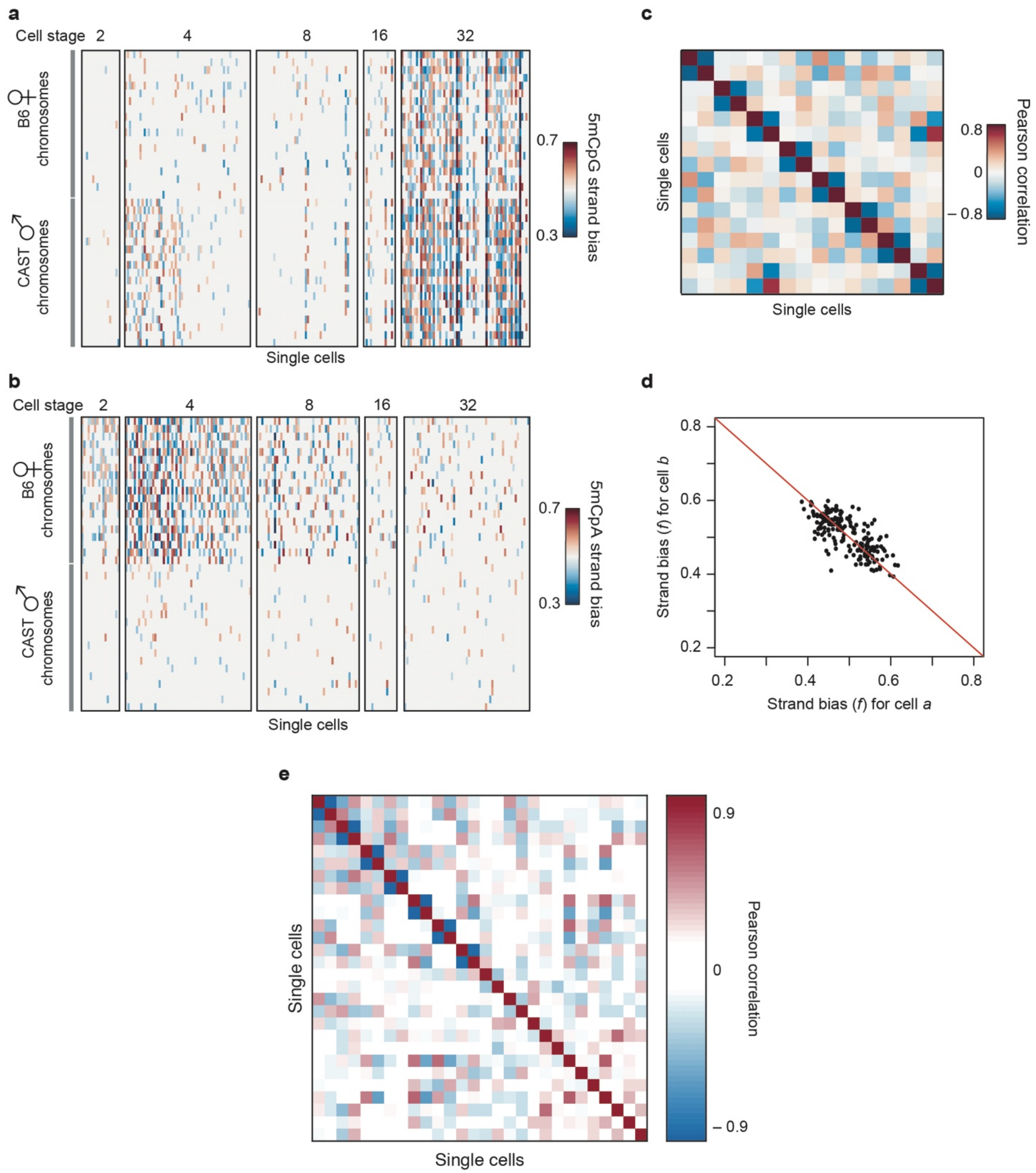


**Supplementary Figure 2 | DNA demethylation dynamics in preimplantation mouse embryos.** (**a**) Heatmap shows 5mCpG strand bias for all maternal and paternal chromosomes from the 2- to 32-cell stage of development. The data shows a dramatic increase in 5mCpG strand bias from the 16- to 32-cell stage of development. (**b**) Heatmap shows 5mCpA strand bias for all maternal and paternal chromosomes from the 2- to 32-cell stage of development. For a majority of cells at the 2- and 4-cell stage, the maternal genome displays 5mCpA strand bias that deviates from 0.5. (**c**) Heatmap shows Pearson correlation for the maternal 5mCpA strand bias between pairs of cells at the 2-cell stage of development. (**d**) Pairs of cells in **c** that display strongly anticorrelated 5mCpA strand bias are shown here, suggesting that we can use this method to identify sister cells at the 2-cell stage of development. (**e**) Heatmap shows Pearson correlation for the paternal 5mCpG strand bias between pairs of cells (within the bimodal strand bias distribution) at the 32-cell stage of development. Strongly negative Pearson correlations indicate that we can identify sister cells within 32-cell stage embryos.

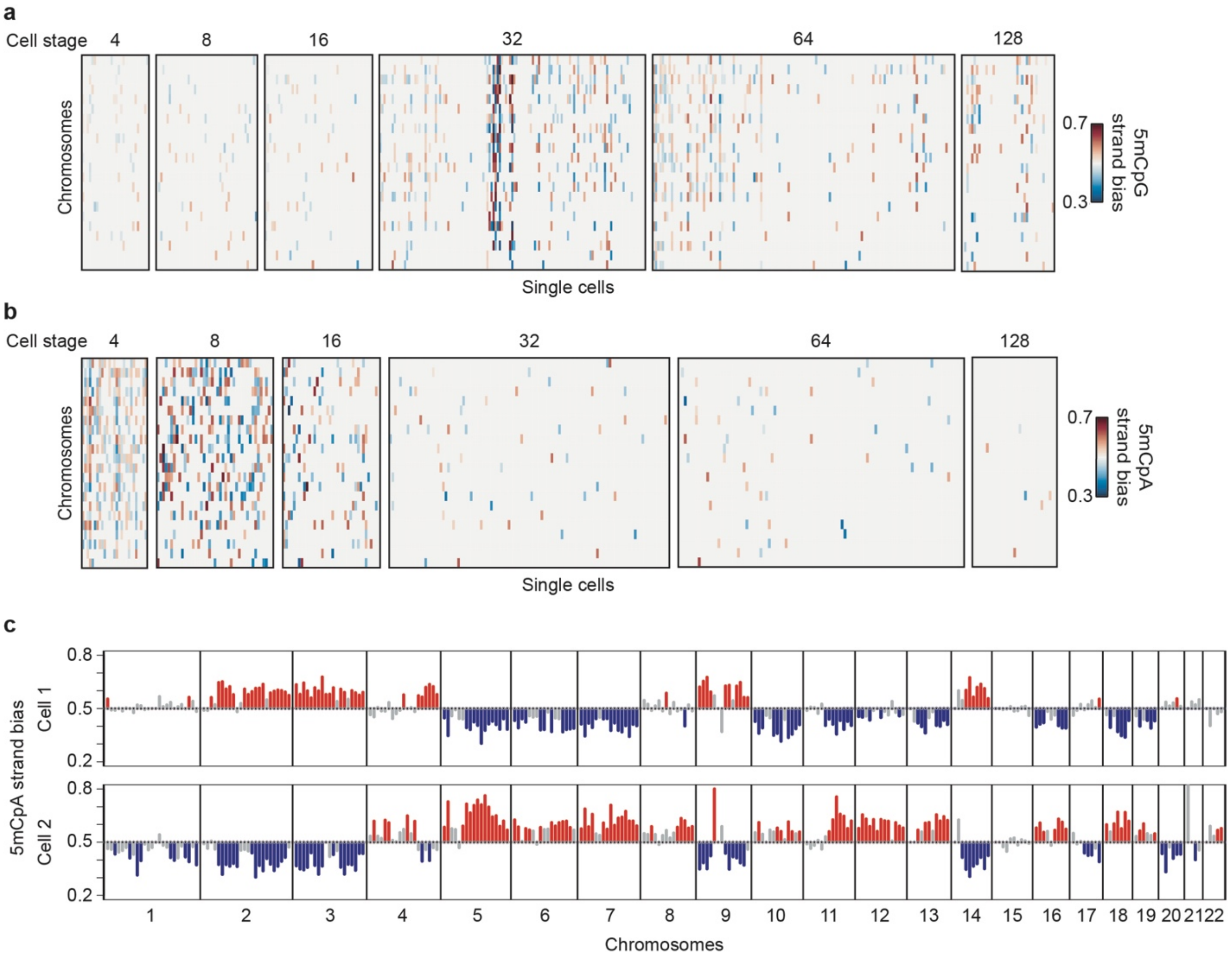


**Supplementary Figure 3 | DNA demethylation dynamics in preimplantation human embryos.** (**a**) Heatmap shows 5mCpG strand bias for all chromosomes from the 4- to 128-cell stage of human development. (**b**) Heatmap shows 5mCpA strand bias for all chromosomes from the 4- to 128-cell stage of human development. 5mCpA strand bias deviates from 0.5 for a large number of chromosomes till the 16-cell stage. (**c**) An example of a pair of cells that display strongly anti-correlated 5mCpA strand bias along the entire genome.

**Supplementary Table 1 | Table listing 8 bp cell-specific barcodes used in scMspJI-seq**

| Cell barcode | Sequence |
|---|---|
| 1 | GCGAGATT |
| 2 | CATTCCAC |
| 3 | CCGATGAT |
| 4 | AGCTTAGC |
| 5 | CGTTACTG |
| 6 | TTCGCTTG |
| 7 | CTACTGCT |
| 8 | AAGAGAGC |
| 9 | AACTGTGG |
| 10 | CACATCAG |
| 11 | AGTCAGTC |
| 12 | CGTTGTCA |
| 13 | TAGGAACG |
| 14 | CCTGATCT |
| 15 | AACGAGCA |
| 16 | GCAGTAAC |
| 17 | TTCTCGAC |
| 18 | GTCCAATC |
| 19 | AACTCACC |
| 20 | CTGCGAAT |
| 21 | ACGTTACC |
| 22 | AGTTGCAC |
| 23 | AATAGCCG |
| 24 | ACCTCTAC |
| 25 | TTCGACGT |
| 26 | TTGATCCG |
| 27 | GTACAGGT |
| 28 | ACCACCTT |
| 29 | GGATTCGA |
| 30 | CCGTTAAG |
| 31 | GTTCGGAA |
| 32 | CCACCATT |
| 33 | CGATCGAT |
| 34 | GTCTGTAC |
| 35 | ACGCCTTA |

| | |
|---|---|
| 36 | CAGGATTC |
| 37 | GGAAGATC |
| 38 | TCAGACGA |
| 39 | TGCGCTAA |
| 40 | GAGAATGC |
| 41 | TTAGCGTG |
| 42 | TGAAGGCT |
| 43 | CTTAGCAG |
| 44 | AAGCTACC |
| 45 | ACATCTGC |
| 46 | CGCATTAC |
| 47 | CCTAGATC |
| 48 | CATCCAGA |
| 49 | GGTCTTGA |
| 50 | GACAGATG |
| 51 | GAACAGCT |
| 52 | ACGAGCAA |
| 53 | TCCTTCTC |
| 54 | GCGTGTAA |
| 55 | CAGCCATA |
| 56 | GAATTGCC |
| 57 | AATCAGCC |
| 58 | CTCAACAC |
| 59 | GCAGATAC |
| 60 | TCGCTTGT |
| 61 | AGTCTTCG |
| 62 | TAGAGGCA |
| 63 | CCTTGGTT |
| 64 | AGAACGCA |
| 65 | GTATACGC |
| 66 | ACTGCTAG |
| 67 | ATCGGTGA |
| 68 | GACCATGA |
| 69 | TCCAAGGT |
| 70 | GCCAACAT |
| 71 | GCGTCAAT |
| 72 | AGCCAAGT |
| 73 | ACGTCAGA |
| 74 | TCACCTGA |

| | |
|---|---|
| 75 | GCAATCCT |
| 76 | AATTCGCC |
| 77 | TGAAGCTC |
| 78 | GTCCGATA |
| 79 | CCTGTAGT |
| 80 | CAGACTGT |
| 81 | TGTAGCCT |
| 82 | GATGCCAT |
| 83 | AACGGCAT |
| 84 | GATAGCAC |
| 85 | TACGGTTC |
| 86 | TGGTTGGA |
| 87 | TCGTGTAC |
| 88 | TAGCGGAA |
| 89 | CTAGGCTA |
| 90 | GCTGTGTA |
| 91 | CAGGTCTT |
| 92 | AAGAGCCA |
| 93 | GCATGACT |
| 94 | TTACGGTC |
| 95 | ACGCATAC |
| 96 | GATGCAAC |
